# High-Resolution Spatial Proteomics Characterises Colorectal Cancer Consensus Molecular Subtypes

**DOI:** 10.1101/2025.05.07.652478

**Authors:** Batuhan Kisakol, Anna Sturrock, Sanghee Cho, Mohammadreza Azimi, Andreas U. Lindner, Manuela Salvucci, Elizabeth McDonough, Joanna Fay, Tony O’Grady, Niamh McCawley, John P. Burke, Deborah A. McNamara, John Graf, Simon McDade, Daniel B. Longley, Fiona Ginty, Jochen H.M. Prehn

**Affiliations:** Department of Physiology and Medical Physics & RCSI Centre for Systems Medicine, RCSI University of Medicine and Health Sciences, 123 St Stephen’s Green, Dublin 2, Ireland; GE HealthCare Technology & Innovation Center (Formerly GE Research Center), 1 Research Circle, Niskayuna, NY, 12309, U.S.A; Institute of Microbiology and Virology, Rīga Stradiņš University, Riga, Latvia; Department of Pathology, Beaumont Hospital, Dublin 9, Ireland; Department of Colorectal Surgery, Beaumont Hospital, Dublin 9, Ireland; Queens University, School of Medicine, Dentistry and Biomedical Sciences. Patrick G Johnston Centre for Cancer Research, 97 Lisburn Road, Belfast BT9 7AE, United Kingdom of Great Britain and Northern Ireland

## Abstract

**Background:** Identification of the consensus molecular subtypes (CMS) opened significant potential for understanding the tumor biology and intertumoral heterogeneity of colorectal cancer (CRC). However, molecular subtyping in CRC traditionally relies on bulk transcriptomics, therefore, lacks spatial and single-cell level aspect.

**Methods:** We constructed tissue microarrays using tumor cores from 222 CRC patients. Arrays were stained and imaged using 54 cell identity and cancer hallmark markers, delivering spatially resolved protein profiles of >2 million cells. RNA sequencing data and CMS classification were also available for these patients. After segmentation of cancer, stromal and immune cells, we investigated intratumoral heterogeneity within CMS subtypes using spatially resolved single-cell protein profiling (>2 million cells). We compared cell types, their spatial organization and their expression of cancer hallmark-related proteins in CMS 1-4 subtypes.

**Results:** We revealed tissue atlases illustrating the cell types/states, spatial heterogeneity, cellular neighborhoods, cellular network, and single-cell protein profiles of CMS tumors. CMS1 tumors had more CD3^+^, CD8^+^, and PD1^+^ immune cells that were found in the epithelial layer frequently. CMS1 was also associated with higher levels of metabolic reprogramming markers such as upregulated glycolysis. CMS2 showed immune segregation, reactive stroma patterns and higher levels of apoptotic and proliferative signaling proteins. CMS3 exhibited clustered cancer cells with high RIP3 levels, suggesting a pro-inflammatory microenvironment. CMS4 displayed stromal-centric and immune-evasive tumors characterized by decreased HLA-1 levels and regulatory T-cell exclusion from epithelium.

**Conclusion:** We present a spatial protein atlas of CRC at single-cell resolution and demonstrate novel aspects of CMS tumour structures.

## Introduction

Colorectal cancer (CRC) poses a huge worldwide health threat as it ranks second in cancer-related deaths and represents the third most frequently diagnosed cancer ^1^. Even though the incidence and mortality rates have improved in the last decades with the emergence of personalised medicine approaches, tumour heterogeneity continues to present challenges for clinical treatment ^2,3^. The proposition of consensus molecular subtypes (CMSs) by Guinney *et al*, ^4^ unlocked vast potential for understanding and therapeutic targeting of intertumour heterogeneity in CRC. While CMS offers a valuable understanding of tumour biology, its dependence on bulk transcriptomic data limits its ability to discern intratumour heterogeneity precisely. Cell-cell interactions and spatial characterisation of the tumours that contribute to CRC heterogeneity also cannot be identified.

Due to these limitations of bulk transcriptomics, recent studies focused on single-cell or spatial transcriptomics to identify cell-level patterns in CMS subtypes ^5–7^. Collectively, these studies have shown how tumour, stromal, and immune landscapes and cell states within these landscapes differ in the different CMS subtypes. Chowdhury *et al*. also developed single-cell CMS (scCMS) assignments and demonstrated the cell compositions of CMS tumours containing different types of scCMS using scRNA-Seq samples from 10 patients ^7^. Of note, the study demonstrated that individual CRC tumours may be composed of tumour cells displaying different CMS traits. Recently, using 14 CRC patient samples, Valdeolivas *et al*. found that spatial and transcriptomic patterns of CMS can potentially lead to the discovery of novel therapeutic targets^8^.

However, the spatial organisation of different cell types within different CMS tumours and their cell ‘state’ in relation to their proximity to other cell types has yet to be explored systematically. Moreover, many spatial or scRNA studies to date have been limited to small sample sizes. In this study, we used multiplexed immunofluorescence and analysed more than 2 million cells in 632 tissue microarrays (TMA) from 222 CRC patients to expand our knowledge of CMS tumours’ protein profiles, heterogeneity, and spatial characteristics. We utilised spatially resolved single-cell level expressions of 54 proteins, ranging from cell proliferation, differentiation, and extrinsic and intrinsic apoptosis pathway markers to metabolic and immune cell markers, to create a cell atlas for CMS tumours. Our study offers an improved understanding of CMS subtypes by providing a comprehensive overview of subtypes with spatial context at the single-cell level, integrating protein and cell type information.

## Materials and Methods

### Patient Cohorts

Formalin-fixed, paraffin-embedded primary tumour tissue sections were obtained from two different cohorts. Colorectal cancer tumours are collected from Beaumont Hospital (RCSI, Ireland), Cohort 1 and Queen’s University Belfast (QUB, UK), Cohort 2. A clinical summary of both cohorts can be found in Supplementary Table 1. For cohort 1, tissues were provided by the Beaumont Hospital Colorectal Cancer Biobank with written consent provided by all patients. Institutional ethical approval was granted by the Beaumont Hospital Research and Ethics Committee (References 08/62 and 19/46). For cohort 2, tissues were supplied by the QUB Department of Pathology with written consent provided by all patients and institutional ethical approval granted (NIB12-0034).

### Transcriptomics and CMS classification

Processing the bulk transcriptomics was performed separately for both cohorts. Cohort 1 samples include four different sub-cohorts. All subcohorts were processed with Illumina HiSeq 4000 and processing steps included removing optical duplicates and cleaning the adaptors. The first two sub-cohorts were sequenced using the KAPA RNA-Seq kit, mapped with STAR (RRID:SCR_004463) ^9^, and reads were generated with FeatureCounts (RRID:SCR_012919) ^10^. In the last two sub-cohorts, Lexogen RNA-Seq kit was used, and reads were mapped with HiSat (RRID:SCR_015530) ^11^. All the read counts were separately processed with DESeq2 (RRID:SCR_015687) ^12^ and applied “rlog” transformation. The detailed processing of the CRC transcriptomics samples for Cohort 2 was previously described by Allen *et al* ^13^. In both cohorts, after the gene expression counts were calculated, CMSclassifier ^4^ tool was used to predict CMS classes from microarray or RNA-seq counts.

### Multiplexed Imaging Workflow

For the MxIF study, TMAs from FFPE tissue blocks 1-mm-diameter cores were produced (three TMAs per patient on average). The protocols for the MxIF processes were described in Gerdes *et al*. ^14^ using Cell DIVE™. MxIF staining and imaging were performed as previously described ^15–17^. On average, TMAs had three cores from each tumour. All the antibodies are shown in Supplementary Table 2. After imaging, each core underwent quality control steps explained previously in Cho *et al*.^*16*^ These filtering steps include: These filtering steps include: 1) For epithelial cells, the number of nuclei for each cell to be 1-2, the area of nuclei/cytoplasm/membrane to be bigger than 10 and lower than 1500 pixels, as well as the total area to be bigger than 50 and lower than 3500 pixels. 2) for stromal cells, the total area (only nuclei) to be larger than 30 and lower than 3500 pixels, and lastly, 3) for both stromal and epithelial cells, a good alignment score (>80%) for staining compared to the first round. Following QC, the spatial single-cell protein expression data were normalised to reduce batch effects and transformed using log2 transformation following the procedures described by Graf *et al* ^*18*^.

### Immune Cell Classification

CD3, CD4, CD8, CD20, FOXP3, and PD1 markers were used to identify immune cells. As previously described by Stachtea *et al*. ^*15*^, a “Cell Auto Training” ^19^ model was trained on manually annotated immune cells, and then this model was used to identify immune cells in all tumour cores. After this step, each cell was attributed with a binary class depending on the immune cell positive and epithelial markers (PCK26 and DAPI). The decision tree assignment of leukocytes based on these attributes is shown in Supplementary Figure 1.

### Spatial Analysis

Following immune cell classification, proximity and network analyses were conducted using Python (v3.10, RRID:SCR_008394). The process comprised several steps and is illustrated in Figure 1E:

**Figure 1:**
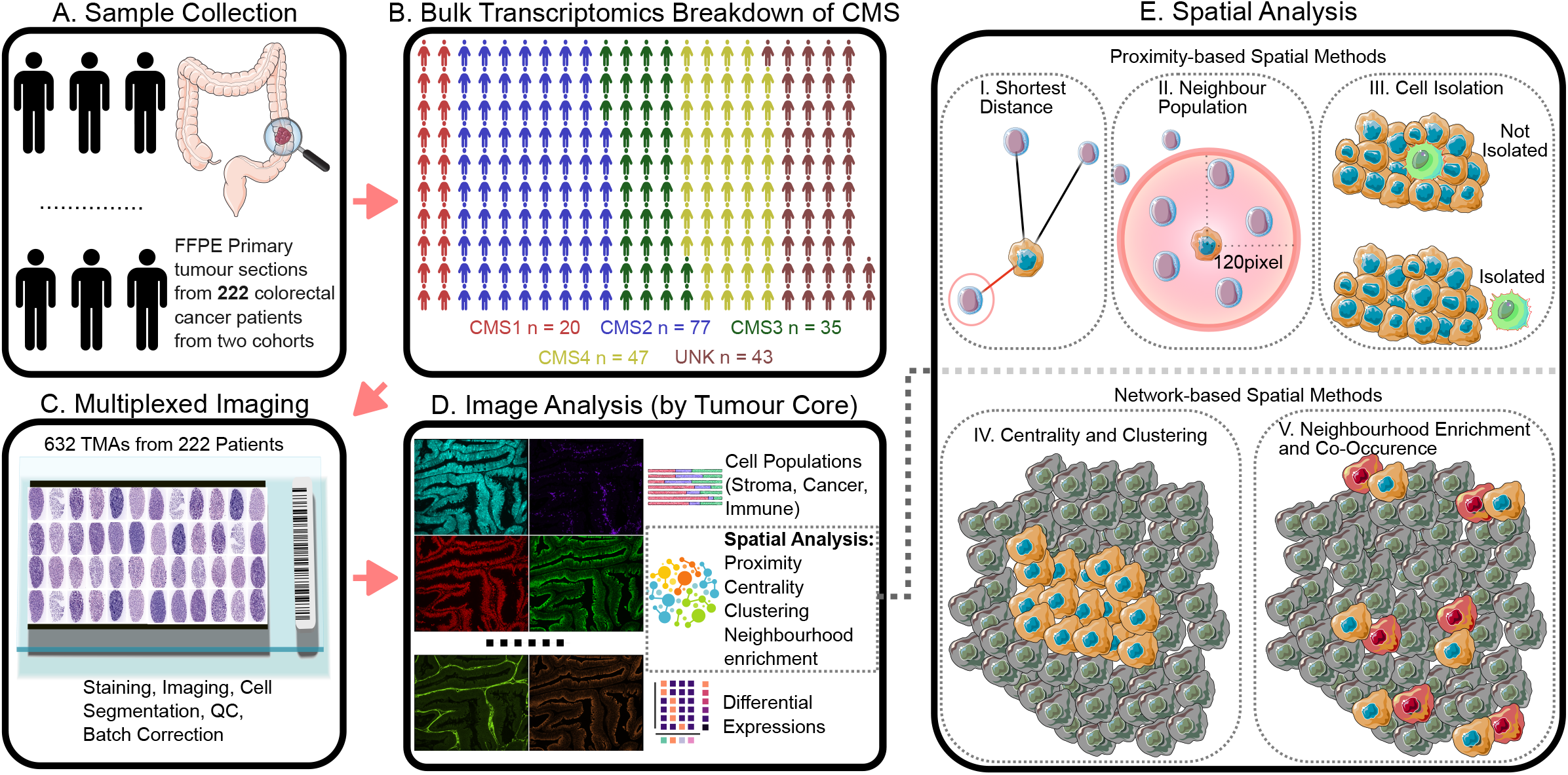
Workflow of the study. Sample collection (A) is followed by bulk transcriptomics to identify CMS subtypes (B). Tumour slides from each core were selected and each TMA analysed with the Cell Dive MxIF technology, staining the TMAs with 54 protein markers (C). Lastly, downstream analysis of the multiplexed images included calculating cell populations, finding spatial patterns within tumour microenvironments, and differential expression analysis of the 54 proteins (D). Then the scheme demonstration of the spatial image analysis methods employed (E). (i) Image demonstrating the analysis of the minimum distance between cell types. (ii) Scheme showing cell number counts (for each cell type) surrounding individual cells. (iii) Analysis of the spatial position of each cell relative to stromal or cancer cell clusters. For this method, stromal or cancer clusters were identified with the K-Means Clustering method based on the spatial location of the cells. Using the Convex Hull Method, the borders of each cluster were then determined. Finally, a binary score was assigned to each cell based on whether or not they were inside the cancer or stromal clusters. (iv) Scheme demonstrating global centrality score analysis. This method was employed to determine whether certain cell types were clustered together or located in central positions compared to the rest of the cells. (v) Network-based neighbourhood enrichment and co-occurrence show whether the cell type pairs were randomly distributed or co-occurred.

### Proximity analysis

The shortest distance between different cell types (e.g., cancer to immune, immune to stroma) was calculated for each cell. These distances were then averaged to represent each tumour core.

### Neighbourhood population

For each cell, the number of different cell types within a 120-pixel/39µm radius was counted. These counts were subsequently averaged for each tumour core.

### Cell isolation analysis

K-means clustering, based on spatial location, was applied to stromal and cancer cells within each tumour core. The borders of each cancer or stromal cluster were determined using the convex hull method. Average isolation scores for each cell type in each tumour core were generated by averaging these results.

### Network analysis

The Squidpy^20^ library was utilised to create network graphs based on cell locations. Each cell was connected to its five nearest neighbours, resulting in cell-level connection networks. Centrality scores, average clustering coefficients, neighbourhood enrichment, and co-occurrence scores were then calculated for each tumour core. Two variations of the network analysis were performed: one treating all immune cells as a single ‘immune’ cluster, and another using specific immune cell classifications (helper, cytotoxic, regulatory, DN T cells, B cells, and other immune cells). Moran’s I scores were generated using the Squidpy library as well.

### Protein Expression Comparison (Differential Expression)

Protein expression comparisons across CMS were conducted following a multi-step process. First, protein expression data were aggregated for each tumour, with cancer, stromal, and immune cells analysed separately. The median expression value was then calculated to represent each protein at the tumour core level. Differentially expressed proteins among the CMS groups are illustrated in Figure 4A-B.

### Statistical Analysis

For statistical analysis, the Mann-Whitney U test was employed to compare each CMS against the combined remainder (e.g., CMS1 versus others, CMS2 versus others, etc.). This non-parametric test was used under the assumption that samples were randomly drawn, independent, and measured on at least an ordinal scale, with similar distribution shapes across groups. For each comparison, the U statistic, sample sizes, exact p-values, and effect sizes were calculated.

For comparisons across multiple CMS groups, the Kruskal-Wallis H test was used to assess differences in distributions, assuming independence between groups and ordinal data.

To assess the relationship between protein expression levels in certain cell types and their distance to other cell types, Spearman’s rank correlation test was used. This analysis assumed paired observations, a monotonic relationship, and at least ordinal-scale data without significant outliers. Reported statistics include the Spearman correlation coefficient, degrees of freedom, and p-values.

To account for multiple comparisons, p-values were subsequently adjusted using the Benjamini-Hochberg method, and all the p-values shown in the figures/tables were adjusted. All the statistical analyses were performed using Python (v3.10, RRID:SCR_008394) and R (RRID: SCR_001905) with ‘stats’ (RRID:SCR_025968), and ‘scipy’ (RRID:SCR_008058) packages.

## Results

### Clinical cohort

A total of 222 patients with CRC were included in this study, with CMS classifications determined through bulk transcriptomics. Of these, 102 were categorised as stage 3, 116 as stage 2, and 4 as stage 1. Regarding sample location, 105 (49%) were from the right-sided colon, 39 (18%) from the left-sided colon, and 69 (32%) from the rectum. 81% of the samples were classified as moderately differentiated, while 39.7% received adjuvant chemotherapy. The most prevalent CMS classification was CMS2, accounting for 77 (34.68%) of the samples. This was followed by CMS4 at 47 (21.17%), unknown/unclassified samples at 43 (19.36%), CMS3 at 35 (15.76%), and CMS1 at 20 (9%). Supplementary Table 1 shows a detailed summary of the patients with the distribution by CMS.

To investigate the cancer, stromal, and immune landscape of colorectal cancer (CRC) tissues at the spatial single-cell level, we used 632 tumour cores that were derived from these 222 CRC patients at resection and performed Cell DIVE™ multiplexing of 54 proteins in the primary tumours (Figure 1, Methods). Following image processing, segmentation, and quality control ^14,17^, a total of 2,050,263 cells were analysed.

### Immune cell infiltration patterns across CMS subtypes verify higher infiltration of CD3^+^, CD4^+^, CD8^+^, and PD1^+^ cells in the epithelial layer in CMS1 tumours

We used pan-cytokeratin antibodies (PCK26 and AE1) and nuclear-specific DAPI fluorescent stain to create epithelial and stromal masks as demonstrated previously ^14,17^. Then we classified immune cells based on CD3, CD4, CD8, CD20, and FOXP3 cell-level expression inside the two masks separately, as described by Stachtea *et al*. ^15^. Based on these immune cell masks, we identified cytotoxic T cells (CD3^+^CD8^+^), helper T cells (CD3^+^CD4^+^FOXP3^-^), regulatory T cells (CD3^+^CD4^+^FOXP3^+^), B-cells (CD20^+^) and other leukocytes (CD3^-^CD20^-^CD4^+^ or CD8^+^) (Supplementary Figure 1). We also detected ‘double negative’ (DN) T cells (CD3^+^CD4^-^CD8^-^) which are believed to be mature T lymphocytes that lack CD4 and CD8 co-receptors and function in immune regulation, cytotoxicity, and cytokine production^21^.

As shown in Figure 2A, we observed higher immune cell infiltration in CMS1 and higher cancer cell abundance in CMS2. Specifically, CMS1 tumours contained more CD3^+^, CD4^+^, CD8^+^, and PD1^+^ cells (Supplementary Table 2, p-values<0.01). CMS1 cores had an average 1.78-to-2.08-fold abundance of immune cells compared to cores of CMS2-4 (Figure 2B and Supplementary Table 2). Especially, the population of cytotoxic T cells was significantly higher than in other CMSs with an average of 2.19-to-3.76-fold change. Of note, T and B cells were more likely to be inside the epithelial cell layer in CMS1 when compared to the other CMS subtypes, as demonstrated in Figure 2C. Although CMS1 tumours exhibited a numerically higher immune/cancer cell ratio, this was not found to be statistically significant (p-value = 0.479, Supplementary Table 2). However, we observed a lower CD4^+^/CD8^+^ ratio in CMS1 and a higher ratio in CMS2 compared to the other subtypes (Supplementary Table 2, p-value<0.01). These findings highlight the distinct immune cell landscape within CMS1 tumours, characterised by increased infiltration of CD3^+^, CD4^+^, CD8^+^, and PD1^+^ cells in the epithelial layer.

**Figure 2:**
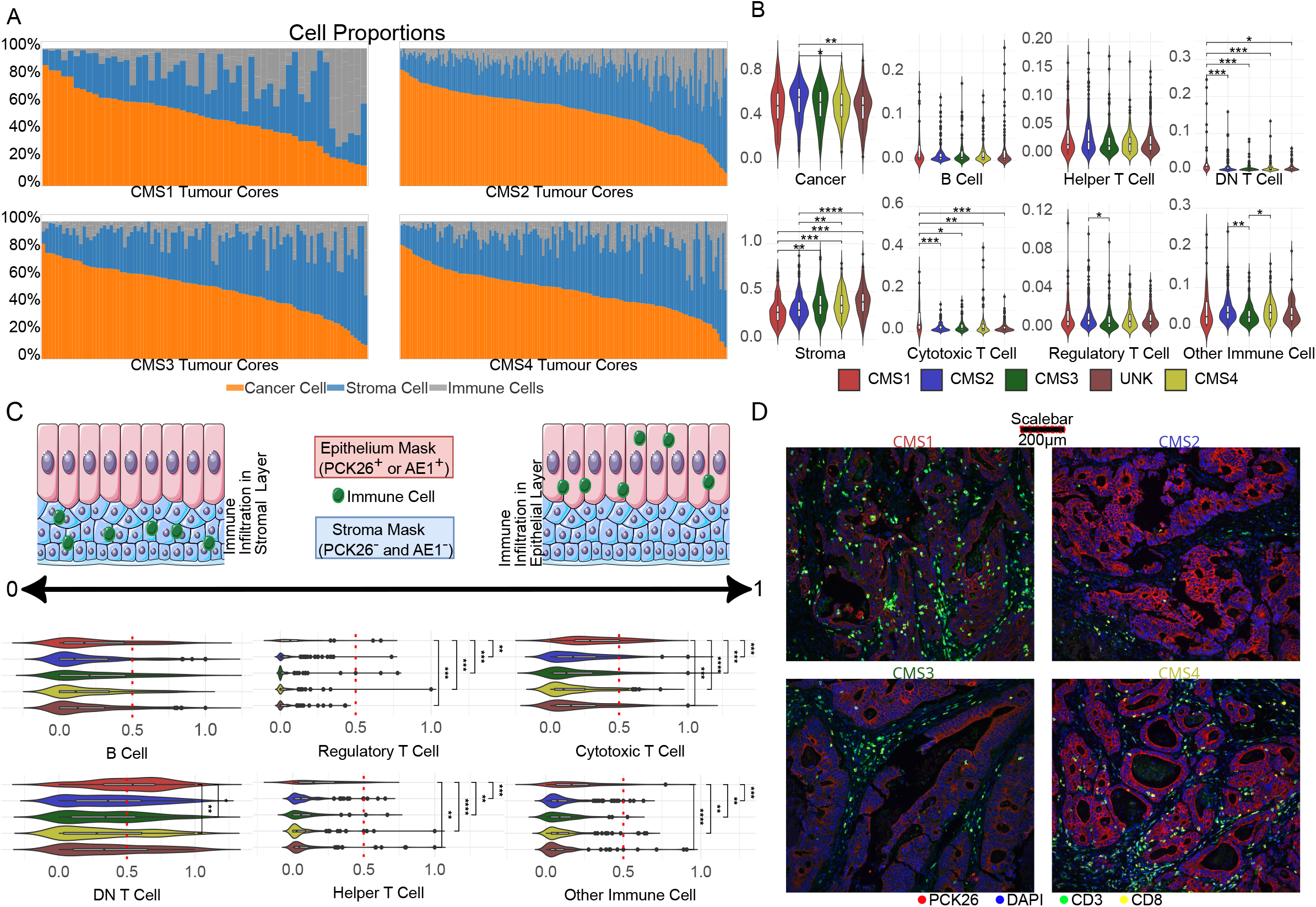
Comparative analysis of tumour microenvironment composition between CMSs shows enriched immune cell infiltration in CMS1. (A) Bar chart representation of the cell type population in each tumour core. (B) Box plot of the cell population for each CMS subtype at tumour core level. (C) Box plot illustration of the immune cell types and the layer they belong to. A score of 0 represents all the immune cells found to be in the stromal layer, and a score of 1 represents all immune cells found in the epithelial layer. Each dot represents a tumour core. (D) MxIF images from each subtype. Representative images were selected by sorting each subtype samples according to their cancer cell abundance, then choosing median or close-to-median samples from each subtype. Colours: DAPI-Blue, PCK26-Red, CD3-Green, and CD8-Yellow. Scale bar = 200 □m.

MxIF images in Figure 2D, selected from samples exhibiting cancer cell abundance proximate to the median within each subtype, provide comparable visual representations. This controlled selection process ensures that the presented images are representative of the central tendency of each group rather than reflecting outliers or randomly chosen specimens. These images visually corroborate the observed trends: CMS1 exhibits elevated immune cell presence within the epithelial compartment, CMS2 demonstrates very little immune cell infiltration, and CMS4 displays pronounced stromal immune cell infiltration

### CMS3 and CMS4 have fewer regulatory T cells in proximity to cancer cells

To understand spatial relationships between cancer cells and immune cells, we analysed the proximity and neighbourhood of these cell types in each tumour core. This analysis was extended to include other stromal cells, i.e. cells that were negative for all epithelial or immune markers (‘non-immune stroma’, hereafter referred to as ‘stroma’), The stromal, tumour and immune cell classification was based on previously published algorithms^14,17^ and is described in Methods. We calculated the distance between the cells in each core and used the nearest distance between cell types to determine proximities between them. We observed that helper T cells, regulatory T cells, cytotoxic T cells, and B cells were closer to cancer cells in CMS1, and that DN T cells were separated from stromal cells (Supplementary Table 3). As demonstrated in Figure 3A, our analysis revealed a significantly reduced average distance between cancer cells and cytotoxic T cells in CMS1 compared to other subtypes. The distance between DN T cells and B cells was also lower in CMS1 (Supplementary Table 3). We also calculated how many immune cells were located within 120 pixels (1 pixel = 0.325 µm) of stromal and cancer cells. We found significantly higher numbers of T cells in proximity to cancer cells and fewer numbers in proximity to stromal cells in CMS1 compared to other subtypes. We also observed that in CMS2, significantly more regulatory T cells were in proximity to cancer cells compared to CMS3 and CMS4 (Supplementary Table 3, p-values < 0.01).

**Figure 3:**
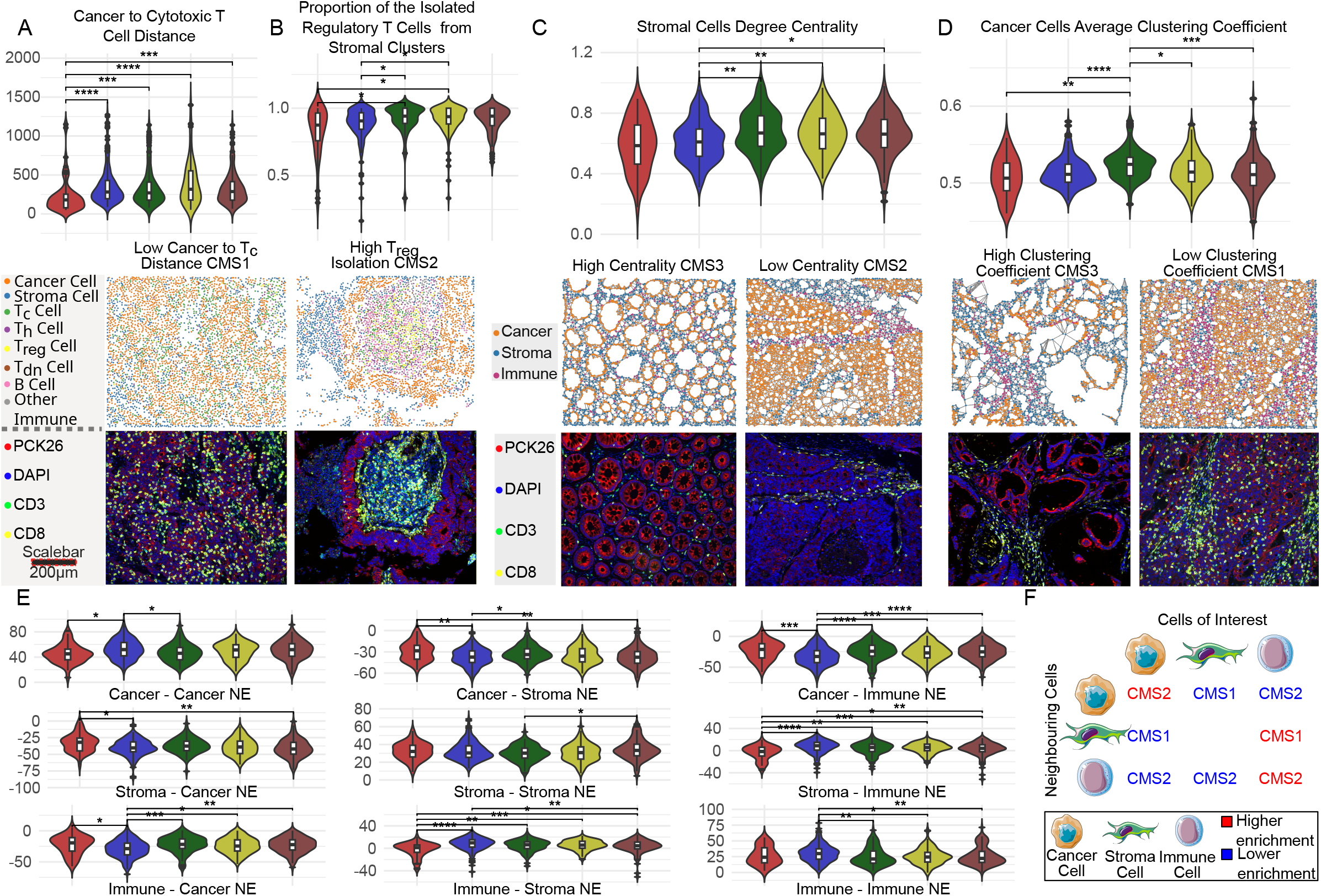
Spatial network analysis reveals low proximity between cancer and cytotoxic T cells in CMS1, high immune-immune and cancer-cancer neighbourhood enrichment in CMS2, and enhanced clustering of cancer cells in CMS3. (A) First row: Boxplot shows the average distance between cancer and cytotoxic T cells for each CMS. Second row: spatial representation of each cell type in an immune-enriched CMS1 tumour. Each dot represents the centre point of a cell, coloured by the cell type. Third Row: MxIF image of the same tumour, colours: DAPI-Blue, PCK26-Red, CD3-Green, and CD8-Yellow. Scale bar = 200 □ m. (B) First Row: Boxplot shows the proportion of Regulatory T cells isolated from stromal clusters for each CMS. Second row: spatial representation of an immune-enriched CMS2 tumour. Third Row: MxIF image of the same tumour, colours: DAPI-Blue, PCK26-Red, CD3-Green, and CD8-Yellow. Scale bar = 200 □ m. (C) First Row: Boxplot shows the stromal centrality scores of each CMS. Second row: representation of the spatial network of cells of a CMS3 tumour with a high stromal centrality and a CMS1 tumour with a low stromal centrality. Third row: MxIF image of the respective tumours, colours: DAPI-Blue, PCK26-Red, and CD3-Green, and CD8-Yellow. (D) First Row: Boxplot illustrates the average clustering coefficient of the cancer cells. Second row: representation of the spatial network of cells of a CMS3 tumour with a high cancer cell clustering coefficient and a CMS1 tumour with a low cancer cell clustering coefficient. Third row: MxIF image of the respective tumours, colours: DAPI-Blue, PCK26-Red, and CD3-Green, and CD8-Yellow. (E) Boxplot representation of neighbourhood enrichment (NE) scores in each subtype. P-values of the pairwise comparisons demonstrated with; *: 0.01 < adjusted p-value <= 0.05, **: 0.001 < adjusted p-value <= 0.01, ***: 0.0001 < adjusted p-value <= 0.001, ****: adjusted p-value <= 0.0001. (E) Summary representation of the (F). It shows the differences in neighbourhood enrichment (NE) scores in each subtype. Rows show the target cells, and columns show the destination cells, demonstrating which neighbourhood pairs were enriched in which CMS subtype (i.e: high immune-immune enrichment in CMS2 and low immune-stromal enrichment in CMS1).

Next, to explore the spatial heterogeneity of immune/stroma/cancer cells, we spatially clustered stromal and cancer cells and determined whether immune cells were found within or outside of these clusters^22^. We categorised immune cells as isolated or non-isolated if they fell inside or outside the borders of stromal or cancer cell spatial clusters. As illustrated in Figure 3B, we found that regulatory T cells were more isolated from the stromal clusters in the CMS1 and CMS2 subtypes. The spatial proximity of regulatory T cells to cancer cells was found to be lower in CMS3 and CMS4 tumours, suggesting a potential lack of immune activation in these subtypes.

### Cellular Network - Global connectivity scores: Cancer cells in CMS3 form tightly interconnected clusters, while stromal cells in CMS3 and CMS4 act as central hubs, suggesting distinct organisational patterns

To investigate the spatial orientation of the tumours, we analysed how the epithelial, stromal, and immune cells were structured within each tumour core by developing a single-cell network model (described in Methods). We generated four global scores for stromal, epithelial, and immune cell groups in every tumour core: closeness centrality, degree centrality, average clustering coefficient (ACC), and connectivity-based neighbourhood enrichment score (see Methods).

We observed that the degree of centrality of stromal cells was higher in CMS3 (0.674 degree centrality) and CMS4 (0.667 degree centrality), suggesting that here stromal cells act as central communication hubs within the tumour microenvironment, interacting with a wide range of neighbouring cells, including cancer cells, immune cells, and other stromal cells, as depicted in Figure 3C.

The average clustering coefficient of cancer cells was high in CMS3 (0.524 ACC). This demonstrated that in CMS3, cancer cells tended to form tightly interconnected clusters or neighbourhoods within the tumour microenvironment. Such clustering behaviour might suggest local coordination or cooperation among cancer cells, possibly facilitating collective migration and invasion within these clusters, as demonstrated in Figure 3D. CMS3 tumours exhibit tightly interconnected cancer cells, while stromal cells in both CMS3 and CMS4 subtypes play a central role in organising the tumour microenvironment.

### Cellular Network - Neighbourhood enrichment: Reduced cancer-immune and cancer-stroma interactions combined with increased immune-immune interactions in CMS2 suggest a unique spatial organisation of the tumour microenvironment

We, then, calculated neighbourhood enrichment scores between cancer, immune, and stromal cells. These scores provide quantitative measures of the degree of connectivity or clustering between different types of cells within a spatial network. Higher scores indicate stronger or more frequent interactions between cells, while lower scores suggest weaker or less frequent interactions.

As illustrated in Figure 3E and Figure 3F, CMS2 tumours showed significant differences in spatial co-localisation of cancer, stromal, and immune cells compared to the rest. First, our analysis revealed a significantly lower enrichment score (ES) of −32.15 on average for cancer cells neighbouring immune cells in CMS2 tumours, indicating a 1.24 to 1.41-fold reduction compared to other subtypes, suggesting a potential lack of interaction between these cell types (Figure 3E, Supplementary Table 3, p-value < 0.01).

This could be a consequence of the immune-desert microenvironment of the CMS2 tumours; however, we also observed high immune-to-immune cell neighbourhood enrichment in CMS2.

The average ES for immune-immune cell interactions was significantly higher in CMS2 tumours (30.14) compared to other subtypes (a 1.13-1.20-fold increase), indicating a greater degree of clustering and potential cooperative behaviour among these cells. Finally, cancer-stroma ES was significantly lower in CMS2 compared to CMS1 and CMS3, with an average of 1.27 and 1.13 fold reduction (Figure 3E, Supplementary Table 3, p-value < 0.01).

Therefore, the observed reduction in cancer-immune and cancer-stroma interactions, coupled with increased immune-immune interactions in CMS2 tumours, indicates a distinct spatial organisation of the tumour microenvironment.

### Protein Level Heterogeneity across CMS Subtypes

As demonstrated in Figure 4A and 4B (and Supplementary Figure 2-3), we observed heterogeneous protein level distributions among CMS subtypes. CMS1 was characterised by elevated levels of PKM2, HTR2B, BCLXL, GRP78, HK2, and KI67 in cancer cells, accompanied by significantly lower CDX2 expression (p-value < 0.01). CMS2 exhibited decreased levels of TIGAR, GLUT1, and ALDH1, while EPCAM, CASP9, and SMAC were upregulated in cancer cells. CMS3 displayed increased RIP3 expression in cancer cells and elevated SDHA and cIAP1 levels in stromal cells. CMS4 was associated with higher β-catenin expression in cancer cells and decreased HLA-1 expression across all cell types (cancer, stromal, and immune). This demonstrated different cell states related to CMS subtypes.

**Figure 4:**
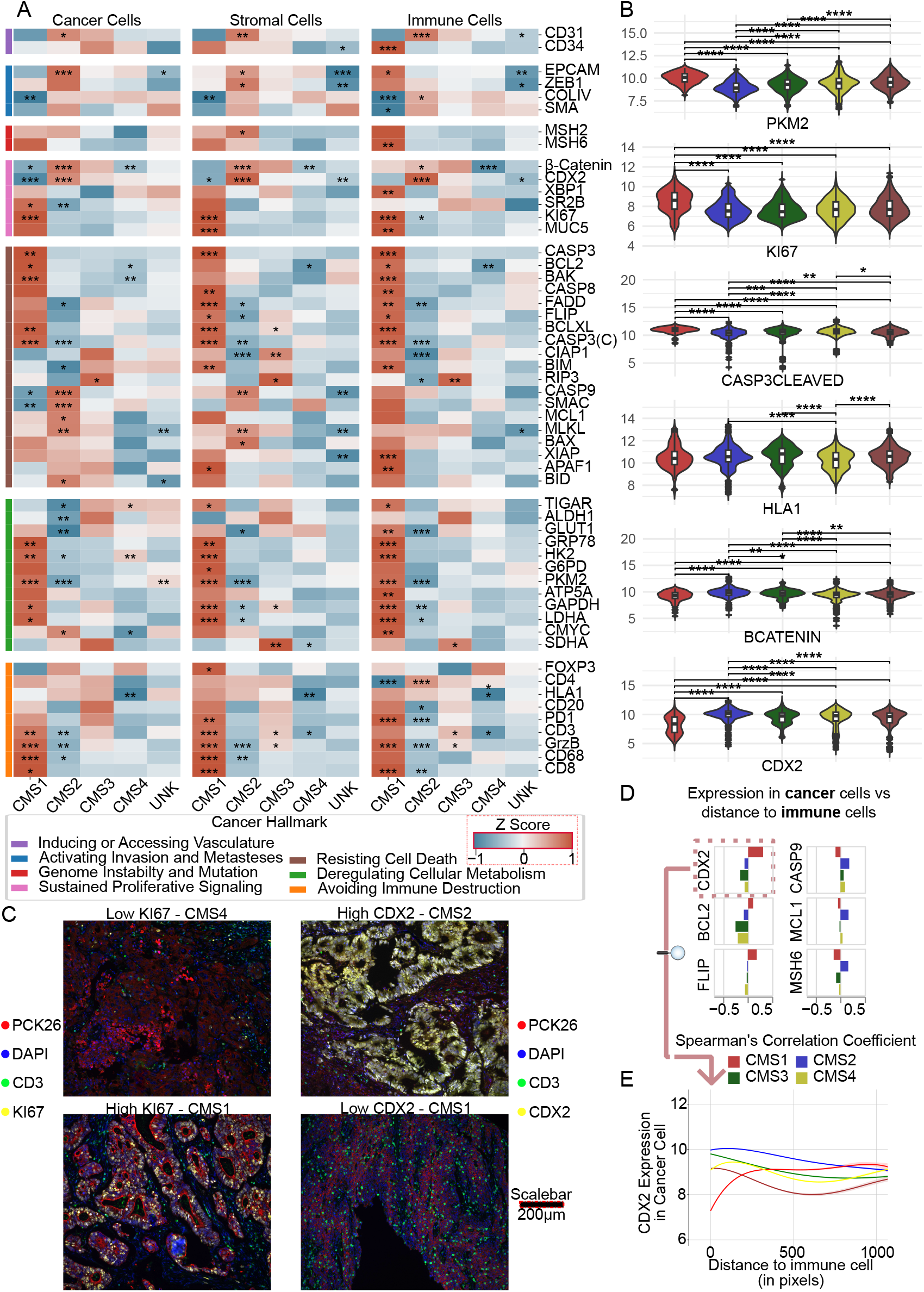
Protein expression profiles of the CMS in cancer, stromal and immune cells show different cancer hallmark patterns. (A) Heatmap representation of the core-level comparison of the protein expression levels in different CMS subtypes. One vs Rest comparison results are demonstrated with; *: 0.01 < adjusted p-value <= 0.05, **: 0.001 < adjusted p-value <= 0.01, ***: 0.0001 < adjusted p-value <= 0.001, ****: adjusted p-value <= 0.0001. (B) Boxplot of the actual expression levels of the most differentially expressed proteins. P-values of the pairwise comparisons demonstrated with; *: 0.01 < adjusted p-value <= 0.05, **: 0.001 < adjusted p-value <= 0.01, ***: 0.0001 < adjusted p-value <= 0.001, ****: adjusted p-value <= 0.0001. (C) MxIF images of CMS samples showing different levels of KI67 and CDX2 expressions. In the first column, the top image shows a low KI67 CMS4 sample, and the bottom one shows high KI67 CMS1 samples. In the second column, the bottom image shows high CDX2 CMS2 sample, whereas the bottom image shows a low CDX2 CMS1 sample. Colours: DAPI-Blue, PCK26-Red, CD3-Green. In the first column, yellow represents KI67, and in the second column, yellow represents CDX2. (D) Barplots show the correlation between the expression of the proteins in cancer/stroma cells and their distance to the nearest immune cell. (E) Line plot of the expression profile of CDX2 in cancer cells relative to the distance to the nearest immune cell.

### Influence of spatial proximity on protein levels

To investigate the influence of spatial proximity on protein levels, we assessed protein levels in cancer cells relative to their distance from the nearest immune or stromal cell. Greater immune cell distances were associated with significantly lower protein levels in cancer cells. Most notably, CDX2 levels, a key regulator of intestinal development and differentiation, were strongly positively correlated with distance to immune cells in CMS1 contrary to the other subtypes (Spearman’s rank correlation coefficient = 0.3, p-value< 0.001) (Figure 4C-D).

Moreover, in CMS1, we observed a positive correlation between FLIP, BCL2 expression, and distance to immune cells, a trend not seen in other subtypes (CMS1 coefficient = 0.1 and 0.18 respectively, p-value< 0.001). Conversely, BAX expression was negatively correlated with distance to immune cells, particularly in CMS1 (coefficient = −0.27, p-value< 0.001). In CMS3, ALDH1, ATP5A, SMA, XIAP, and XBP1 were negatively correlated with distance to immune cells (coefficients between −0.16 and −0.12, Supplementary Figure 4). In CMS4, cancer cells exhibited increased expression of cMYC and CASP8 when located farther from stromal cells. Conversely, in CMS1, stromal cells showed higher expression of EPCAM and BCL-XL when situated farther from cancer cells (coefficients are between 0.11-0.18, Supplementary Figure 4).

We further investigated the protein expression profiles of stromal cells in relation to their proximity to cancer and immune cells. In CMS2, we observed positive correlations between the expression of MCL1, CASP9, cleaved CASP3, MSH2, and MSH6 in stromal cells and their distance to immune cells (coefficients between 0.12-0.18). Conversely, in CMS1, MUC5 expression in stromal cells exhibited a weak positive correlation (0.08) with distance to cancer cells, while a negative correlation was observed in other groups.

### Assessing Spatial Clustering of Protein Expression Using Moran’s I

Finally, we used Moran’s I score to assess the spatial distribution of each protein in each tumour core. Moran’s I score quantifies how similar the protein expression levels are between neighbouring cells or regions. A high score indicates that protein expression levels are clustered, meaning similar protein expression values are located close together. A low score implies a random or dispersed pattern of protein expression, suggesting that protein levels are not significantly correlated with the spatial location of cells. We observed different spatial autocorrelation patterns of signalling proteins between CMS subtypes. APAF1, ATP5A, β-catenin, BIM, CASP3, CASP8, cMYC, MCL1, MSH2, MSH6, RIP3, TIGAR, and ZEB1 proteins showed higher Moran’s I score in CMS2 (Moran’s I score intervals between 0.45 and 0.58, p-values < 0.01, Supplementary Figure 5). These protein expression levels were not randomly distributed but were clustered within specific areas of the tumour. Conversely, cleaved CASP3, GLUT1, and SR2B demonstrated low Moran’s I scores in CMS1, which might imply a more uniform expression of these proteins and possibly a more homogeneous tumour microenvironment (Supplementary Figure 5).

## Discussion

CMS classification has provided valuable insights into CRC heterogeneity, offering a framework for understanding disease biology and guiding therapeutic strategies. However, while CMS has enhanced our understanding of different molecular landscapes in CRC, it needs the spatial context crucial for fully understanding the complex interactions within the tumour microenvironment. To investigate the spatial structure of different CMS tumours, we used Cell DIVE MxIF and generated spatially resolved single-cell level expressions of 54 proteins for 222 CRC tumours. Initially, we conducted a comparative analysis of cell type abundance across different CMS subtypes. Subsequently, we investigated the spatial distribution of these cell types within tumour cores, examining their proximity and neighbourhood relationships. To further investigate spatial context, we constructed cell-cell interaction networks. This enabled the calculation of global connectivity and enrichment scores for each tumour core and cell type, capturing not only the distances but also the overall spatial organisation and intercellular relationships within the tumour microenvironment. We calculated global connectivity and enrichment scores for each tumour core and cell type using four key metrics: centrality, clustering, neighbourhood enrichment, and co-occurrence scores. Integrating these spatial scores with cell-level protein expressions, this study collectively provided a comprehensive view of the spatial dynamics and cellular interactions in the tumour landscape, offering insights beyond simple proximity or differentially expressed protein analyses.

### Spatial organisation of immune cells, metabolic reprogramming, upregulated glycolysis, and a spatial disconnect between helper T cells and stromal components in CMS1

Our first finding supported previous works ^4,23^ showing high immune cell infiltration, specifically cytotoxic T cells, in CMS1. Using single-cell-level information, we also presented that immune cells in CMS1 are more likely to be found in the epithelial layer compared to other CMS subtypes. It has been shown that cytotoxic T cells and regulatory T cells located near the epithelial layer are associated with a better prognosis ^24–26^. Our findings corroborate these observations by demonstrating abundant cytotoxic T cell infiltration in the epithelial layer within CMS1 tumours, a subtype linked to improved survival^4^. Furthermore, we identified sparse regulatory T cell presence in the epithelial layers of CMS4 tumours, but substantial regulatory T cell presence in stromal clusters, which may correlate with the reported worse prognosis in this subtype ^4,27^. We also investigated the stromal clusters rather than cell-to-cell level distances by clustering non-immune stromal cells and measuring the neighbouring immune cells to these clusters. Our findings showed fewer helper T cells neighbouring stromal clusters in the good prognosis groups, i.e. CMS1 and CMS2. Considering this finding in conjunction with the high infiltration of T cells in the epithelial layer, it could indicate that helper T cells are effectively migrating from the stroma into the tumour epithelium, which might be associated with better anti-tumour responses. This migration pattern is further supported by our neighbourhood enrichment and co-occurrence analyses in CMS1, which revealed low immune-stromal enrichment coupled with higher cancer-stroma enrichment. This spatial organisation suggests a dynamic process where helper T cells may be actively moving from stromal regions into cancer-rich areas, potentially enhancing anti-tumour immunity. More isolated helper T cells in the CMS1 and CMS2 stromal cell groups might also suggest that the molecular structure of the stromal cell clusters in these subtypes provides a supportive environment for cytokine secretion. The lower immune-stromal enrichment could indicate that these isolated helper T cells are positioned to maximise their effect, potentially concentrating their cytokine secretion in key areas. However, the significant differences in the protein profiles of CMS1 and CMS2 suggest that the isolated helper T cells and cancer cells might function differently in each context.

In the protein level expression comparison, we found higher levels of cleaved CASP3 and PD1 in CMS1 in all types of cells (cancer, stroma, and immune) compared to CMS2. Both of these markers are associated with T cell exhaustion^28–30^. Moreover, PD1 expression in cancer, stroma, and immune cells was also lower in CMS2 and higher in CMS1. By binding its ligand, PD-L1, PD1 sends inhibitory signals to the T cells and prevents their activation, following cytokine production^31^. Therefore, in CMS1, helper T cells are likely undergoing either apoptosis, as higher intrinsic and extrinsic apoptotic marker expressions in CMS1 also suggest, or exhaustion through high PD1 expression. These helper T cells may be more pro-inflammatory, but become exhausted and gradually have reduced effects and impaired cytokine production. Whereas in CMS2, isolated helper T cells might be less activated. Similarly, Karpinski *et al* ^32^ reported that a subgroup of CMS2 displayed high naive helper T cell enrichment. Considering the immune-desert phenotype of CMS2, this could result in less suppression of helper T cell responses when they do get stimulated. Therefore, therapies that provide strong T cell stimulation for CMS2 tumours have been proposed previously ^33–35^. Regarding CMS1, we also observed high levels of PKM2 and HK2 expressions in cancer cells, which are key enzymes in glycolysis. It has been reported that glycolytic activity enhances PD-L1 expression in tumour cells ^36^. Overall, in CMS1, the upregulation of glycolytic markers (HK2, PKM2) in both stromal and immune cells implies an elevated metabolic demand, potentially supporting increased immune cell activity. Concurrently, the presence of immune checkpoint markers (PD-1) and apoptosis-related proteins (cleaved CASP3, BAK) in these cells suggests an immunosuppressive microenvironment and potential T cell exhaustion. The proliferative marker Ki67 indicates ongoing cellular activity, likely encompassing both tumour and immune cells. Finally, CDX2 expression of cancer cells in CMS1 was lower in the presence of immune cells. CDX2 is an intestinal cell-specific transcription factor associated with differentiation, cell interactions and may suppress tumorigenesis. Poorly differentiated or undifferentiated intestinal tissue is linked to low or lost CDX2 expression ^37,38^. Previous studies identified loss of CDX2 with CMS1 and CMS4 ^39–41^. Moreover, Chewchuk *et al* reported that CDX2 influences homeostasis in intestinal epithelial cells via modulating local immunological responses and controls immune cell infiltration ^42^. Our observation suggests a potential link between immune activity and loss of differentiation markers associated with CDX2 in CMS, and suggests CDX2 as a possible therapeutic target.

### Cell proliferation with metabolic adaptation, immune segregation, and reactive stroma signatures in CMS2

Of note, CMS2 had distinct spatial and molecular patterns from the other CMS subtypes. Firstly, the higher number of cancer cells, coupled with lower cancer-immune neighbourhood enrichment and higher immune-immune neighbourhood enrichment, suggests a tumour microenvironment where cancer cells are relatively isolated from immune cells, while immune cells cluster together. This spatial arrangement could indicate that an immune response is present but potentially ineffective in directly engaging cancer cells.

The decreased expression of TIGAR and GLUT1 suggests altered glucose metabolism. Moreover, decreased ALDH1 coupled with metabolic alterations, indicates reduced stem cell-like properties. Upregulation of EPCAM, CASP9, and SMAC, along with high levels of CD31, β-catenin, CDX2, MLKL, BID, and cMYC, indicates enhanced epithelial characteristics, apoptotic potential, and proliferative capacity. It has been reported that cMYC can promote glycolysis through GLUT1 and HK2 ^43^. However, in CMS2 CRC, we observed an opposite pattern where GLUT1 and HK2 expressions were low, which potentially suggests a deviation from the classic Warburg effect. Combining this unique metabolic profile with reduced ALDH1 and cancer stem cell-like properties, CMS2 cancer cells might represent a more differentiated state with high metabolic plasticity.

The higher Moran’s I scores for various proteins, including those involved in apoptosis, DNA repair, and EMT, suggest spatial clustering of these molecular features. This might indicate that there are potentially more specific cancer clusters within this subtype. Also, the low CIAP1 and high ZEB1 and BAX protein levels in stromal cells might suggest a more reactive or potentially tumour-supportive stroma. Although CMS2 tumours exhibit low immune infiltration, the expression of MCL1, CASP9, cleaved CASP3, MSH2, and MSH6 in stromal cells remains correlated with their proximity to immune cells, suggesting a potential indirect effect of the immune microenvironment on stromal cell phenotype. These signals might create a molecular gradient that influences the expression of apoptosis-related (MCL1, CASP9, cleaved CASP3) and DNA repair (MSH2, MSH6) proteins in stromal cells based on their distance from immune cells.

### Interconnected cancer cells with inflammatory stroma and metabolic adaptation in CMS3

Regarding the clustering of the cancer cells, our observation of CMS3 having tightly interconnected clusters may indicate more direct cell-to-cell signaling between cancer cells and possibly the exchange of pro-tumourigenic signals and growth factors. The elevated RIP3 levels in CMS3 in cancer, stroma and immune cells point towards a pro-inflammatory microenvironment with necrotic activity. RIP3 is a key regulator of necroptosis, a form of programmed cell death ^44^. Moreover, RIP3 has been implicated in metabolic regulation, such as lipid metabolism ^45^, which may explain the metabolic alteration characteristics of CMS3. We also consider that RIP3 upregulation might be a response to cellular stress within the tightly clustered environment. Also, the accompanying low FLIP expression, an inhibitor of apoptosis, reinforces this notion of increased cell death. The metabolic profile of CMS3, marked by high SDHA in stromal cells, indicates active oxidative phosphorylation in the stroma ^46^. This could be a compensatory mechanism to support the metabolic demands of the interconnected cancer cells or a reflection of the inflammatory microenvironment. Moreover, our observation of higher stromal degree centrality of CMS3 and CMS4 suggests that stromal cells play a crucial role in mediating cellular interactions and signalling pathways within the tumour microenvironment.

### Stromal-Centric and Immune-Evasive CMS4 exhibits dynamic microenvironment adaptation

In addition to the high centrality of stromal cells in CMS4, cancer cells in CMS4 harbour fewer regulatory T cells, combined with decreased HLA-1 expression across all cell types, indicating a potential immune evasion strategy. The increased β-catenin expression in cancer cells aligns with EMT and supports invasive behaviour. Low expression of apoptosis regulators (BCL, BAK) and the proliferation marker cMYC suggest a state of cellular quiescence ^47,48^ or stress adaptation. Based on stromal cell presence, Based on stromal cell presence, the dynamic regulation of cMYC and CASP8 in cancer cells may indicate that cancer cells, when less influenced by the stroma, adopt a more aggressive phenotype characterised by increased proliferation and potential apoptotic resistance. Collectively, these findings suggest CMS4 tumours might be capable of evading immune surveillance with a reactive stroma that adapts to immune cell presence, potentially contributing to treatment resistance and poor clinical outcomes.

## Resource availability

## Lead Contact

Further information and requests for resources and reagents should be directed to and will be fulfilled by the lead contact, Jochen H.M Prehn.

## Materials availability

This study did not generate new unique reagents.

## Data and code availability

- Cell Dive multiplexed images generated and analysed in this paper will be shared by the lead contact upon request.
- All original codes can be found at https://github.com/kisakol/Single_Cell_Protein_GE (It is a private repository that can be accessed upon request. It will be made public after publication).
- Any additional information required to re-analyse the data reported in this paper is available from the lead contact upon request.

## Acknowledgements

BK is supported by Research Ireland through the Research Ireland Centre for Research Training in Genomics Data Science under Grant number 18/CRT/6214, EU’s Horizon 2020 research, and innovation programme under the Marie Sklodowska-Curie grant H2020-MSCA-COFUND-2019-945385. JHMP is supported by US-Ireland Tripartite R01 award from Research Ireland and the Health Research Board (16/US/3301). FG is supported by the National Cancer Institute of the National Institutes of Health under award number R01CA208179. DL and SMD were supported by a US-Ireland Tripartite R01 award (NI Partner supported by HSCNI, STL/5715/15).

## Author contributions

Conceptualization, B.K., A.M., S.C., S.M., D.B.L., F.G., and J.H.M.P; Methodology, B.K., A.M., S.C., M.A., A.U.L., M.S., E.M., J.G., and F.G.; Software, B.K.;Writing – Original Draft, B.K.; Writing – Review & Editing, all the authors; Funding Acquisition, S.M., D.B.L., F.G., and J.H.M.P; Resources, E.M., J.F., T.O., N.M., J.P.B., D.A.M.; Data Curation, S.C. and M.S.; Supervision, F.G. and J.H.M.P

## Supplemental information

Document S1. Figures S1–S6 and Table S1-S3

