## Supplementary Tables and Figures for "High-Resolution Spatial Proteomics Characterises Colorectal Cancer Consensus Molecular Subtypes"

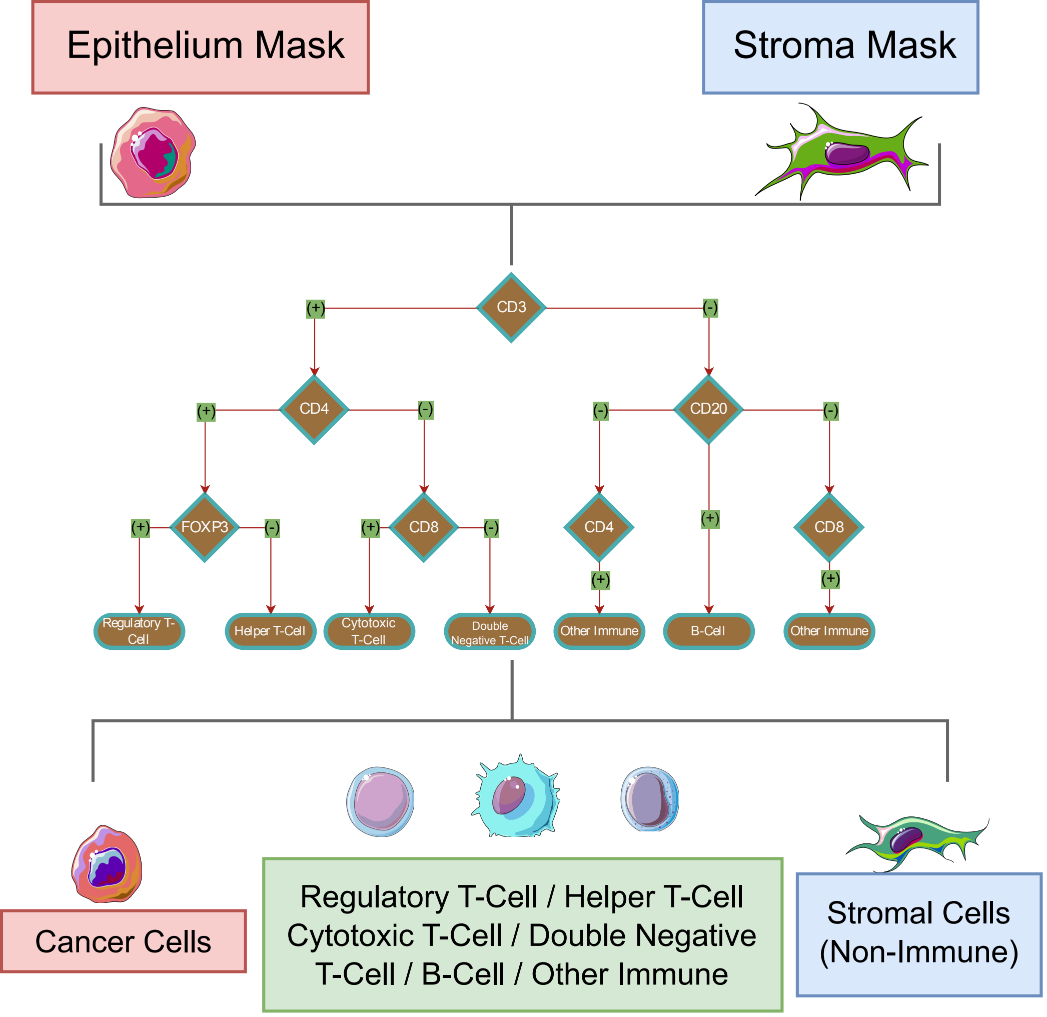


Supplementary Figure 1: Immune cell classification

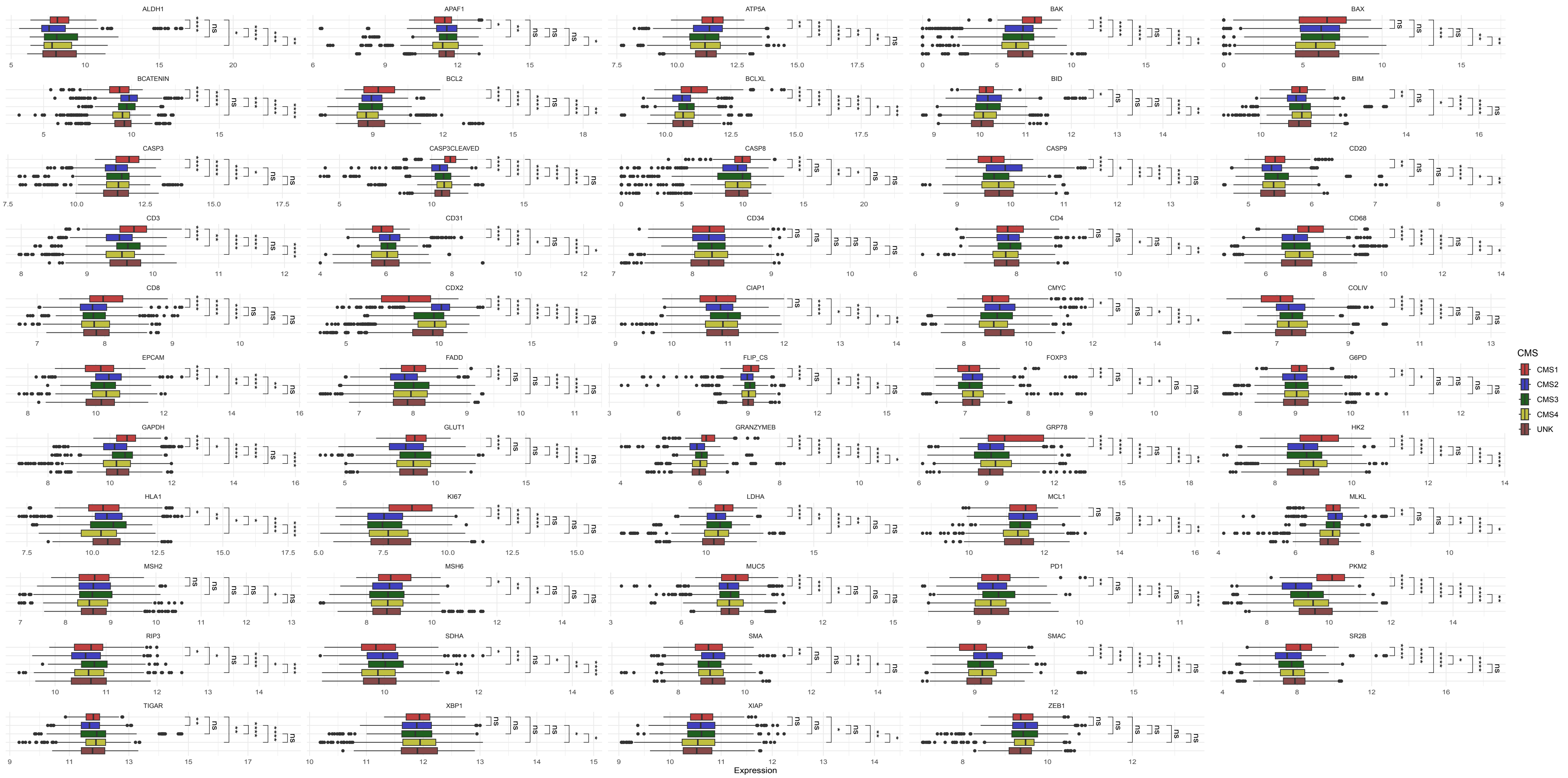


Supplementary Figure 2: Boxplots show the core-level expressions of every protein in cancer cells. Mean values of each protein in cancer cells were calculated for each tumour core. Dots represent tumour cores. P-values of the pairwise comparisons demonstrated with; *: 0.01 < adjusted p-value <= 0.05, **: 0.001 < adjusted p-value <= 0.01, ***: 0.0001 < adjusted p-value <= 0.001, ****: adjusted p-value <= 0.0001.

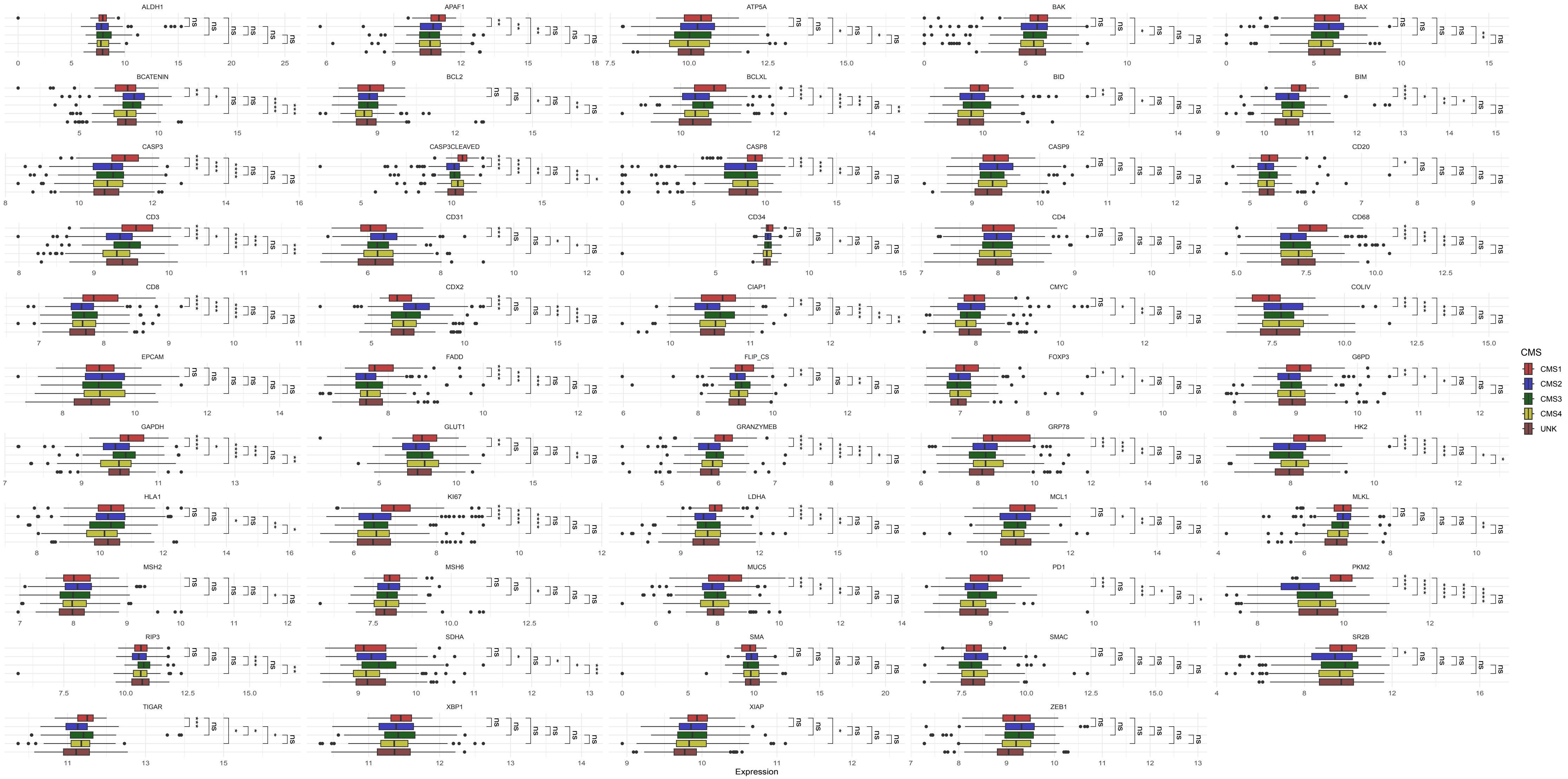


Supplementary Figure 3: Boxplots show thecore-level expressions of every protein in stromal cells. Mean values of each protein in stromal cells were calculated for each tumour core. Dots represent tumour cores. P-values of the pairwise comparisons demonstrated with; *: 0.01 < adjusted p-value <= 0.05, **: 0.001 < adjusted p-value <= 0.01, ***: 0.0001 < adjusted p-value <= 0.001, ****: adjusted p-value <= 0.0001.

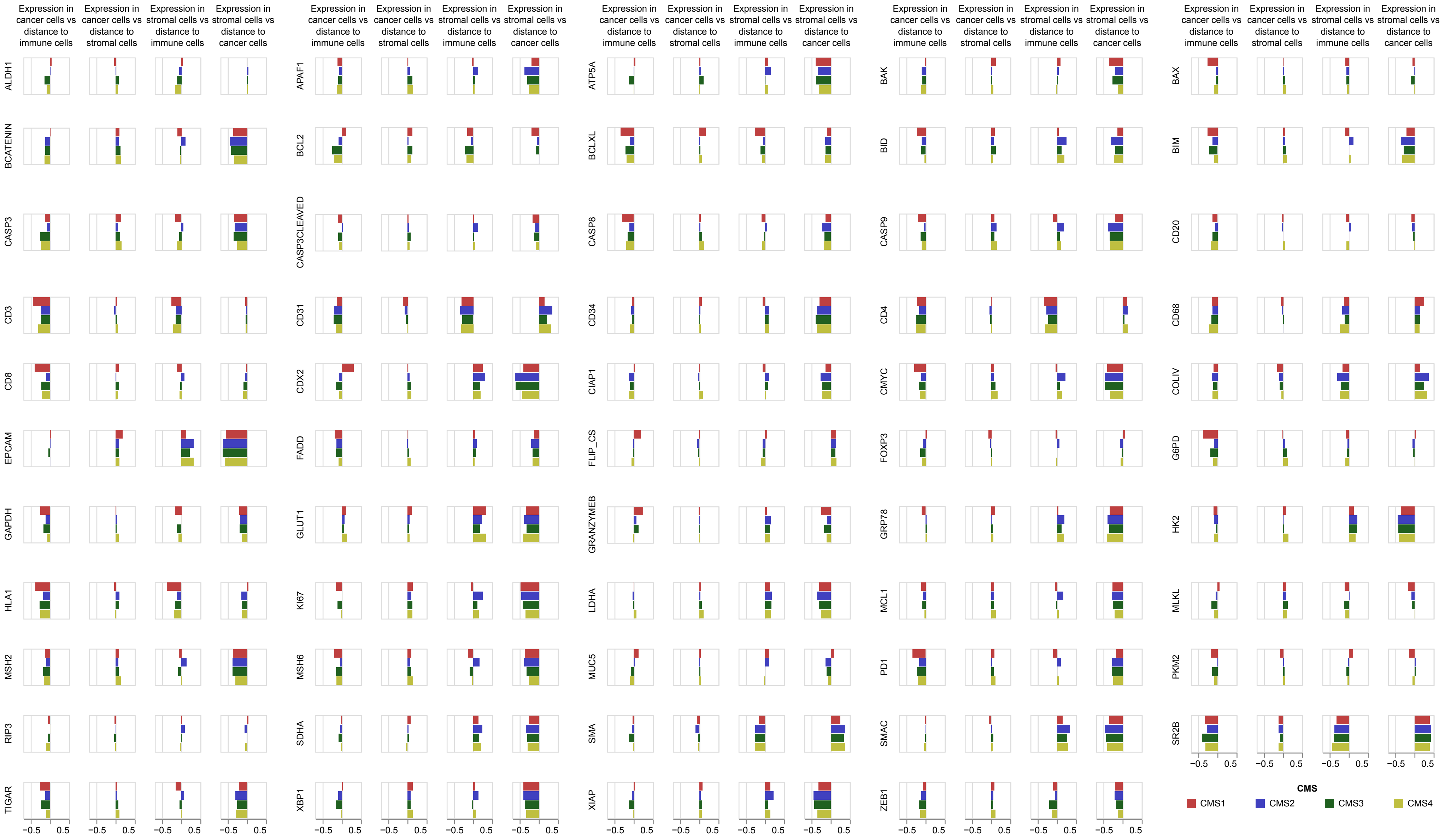


Supplementary Figure 4: Barplots show the correlation between expression of the proteins in the specific cell type and distance to the nearest specific cell type. For each protein, four correlation is shown in the barplots. First row is expression in cancer cells and distance to immune cells. Second row is expression in cancer cells and distance to stromal cells. Third row is expression in stromal cells and distance to immune cells. First row is expression in stromal cells and distance to immune cells. Coefficients were calculated using Spearman's rank correlation.

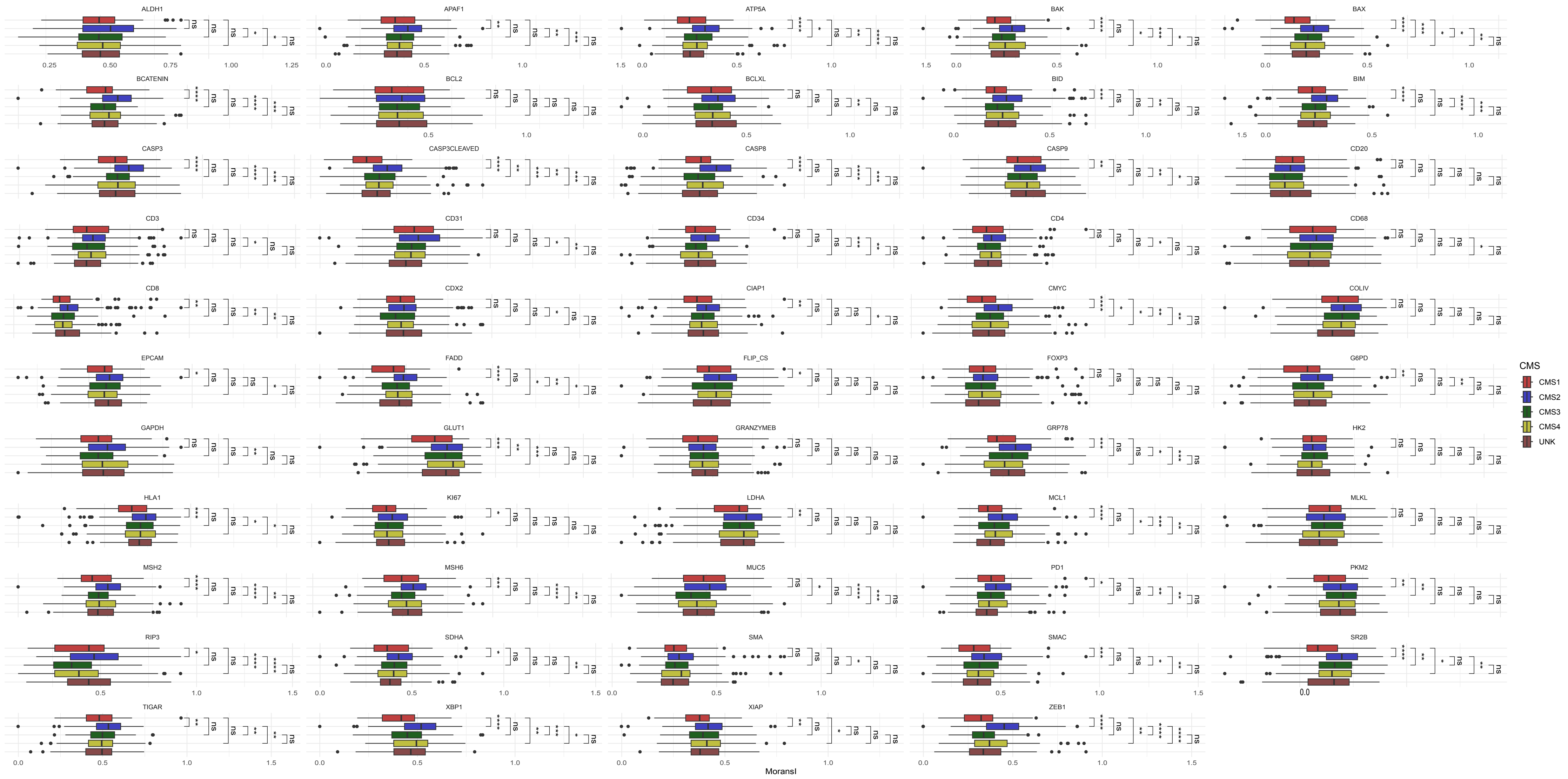


Supplementary Figure 5: Boxplots show the Moran's I score of each protein for every tumour core. Moran's I scores were calculated using only the cancer cells. P-values of the pairwise comparisons demonstrated with; *: 0.01 < adjusted p-value <= 0.05, **: 0.001 < adjusted p-value <= 0.01,  ***: 0.0001 < adjusted p-value <= 0.001, ****: adjusted p-value <= 0.0001.

### Table 1 Samples Summary

| | CMS1 (N=20) | CMS2 (N=77) | CMS3 (N=35) | CMS4 (N=47) | UNK (N=43) | Total (N=222) | p value |
|---|---|---|---|---|---|---|---|
| **Age** | | | | | | | 0.328 |
| N-Miss | 0 | 0 | 0 | 1 | 1 | 2 | |
| Mean (SD) | 73.700 (11.008) | 68.987 (10.169) | 72.657 (13.106) | 70.652 (9.324) | 69.476 (12.381) | 70.441 (11.059) | |
| Median | 75.500 | 71.000 | 74.000 | 70.500 | 71.000 | 72.000 | |
| Q1, Q3 | 69.000, 82.250 | 62.000, 77.000 | 66.000, 83.000 | 65.000, 77.000 | 63.250, 78.000 | 63.000, 78.000 | |
| Range | 44.000 - 89.000 | 37.000 - 88.000 | 38.000 - 94.000 | 51.000 - 87.000 | 46.000 - 88.000 | 37.000 - 94.000 | |
| **Sex** | | | | | | | 0.068 |
| female | 14 (70.0%) | 26 (33.8%) | 12 (34.3%) | 20 (42.6%) | 24 (55.8%) | 96 (43.2%) | |
| male | 6 (30.0%) | 51 (66.2%) | 23 (65.7%) | 27 (57.4%) | 19 (44.2%) | 126 (56.8%) | |
| **stage** | | | | | | | 0.251 |
| 1 | 0 (0.0%) | 2 (2.6%) | 2 (5.7%) | 0 (0.0%) | 0 (0.0%) | 4 (1.8%) | |
| 2 | 12 (60.0%) | 47 (61.0%) | 19 (54.3%) | 19 (40.4%) | 19 (44.2%) | 116 (52.3%) | |
| 3 | 8 (40.0%) | 28 (36.4%) | 14 (40.0%) | 28 (59.6%) | 24 (55.8%) | 102 (45.9%) | |
| **Tumour Location** | | | | | | | 0.545 |
| N-Miss | 0 | 5 | 1 | 3 | 0 | 9 | |
| distal | 3 (15.0%) | 15 (20.8%) | 7 (20.6%) | 6 (13.6%) | 8 (18.6%) | 39 (18.3%) | |
| proximal | 14 (70.0%) | 30 (41.7%) | 15 (44.1%) | 25 (56.8%) | 21 (48.8%) | 105 (49.3%) | |
| rectal | 3 (15.0%) | 27 (37.5%) | 12 (35.3%) | 13 (29.5%) | 14 (32.6%) | 69 (32.4%) | |
| **Differentiation** | | | | | | | 0.084 |
| N-Miss | 0 | 0 | 2 | 0 | 1 | 3 | |
| moderate | 14 (70.0%) | 69 (89.6%) | 26 (78.8%) | 39 (83.0%) | 30 (71.4%) | 178 (81.3%) | |
| poor | 6 (30.0%) | 6 (7.8%) | 3 (9.1%) | 7 (14.9%) | 10 (23.8%) | 32 (14.6%) | |
| well | 0 (0.0%) | 2 (2.6%) | 4 (12.1%) | 1 (2.1%) | 2 (4.8%) | 9 (4.1%) | |
| **Adjuvant Chemotherapy** | | | | | | | 0.545 |
| N-Miss | 0 | 1 | 0 | 1 | 1 | 3 | |
| no | 12 (60.0%) | 46 (60.5%) | 25 (71.4%) | 24 (52.2%) | 25 (59.5%) | 132 (60.3%) | |
| yes | 8 (40.0%) | 30 (39.5%) | 10 (28.6%) | 22 (47.8%) | 17 (40.5%) | 87 (39.7%) | |
| **KRAS Status** | | | | | | | 0.308 |
| N-Miss | 3 | 15 | 6 | 12 | 12 | 48 | |
| mutant | 3 (17.6%) | 19 (30.6%) | 15 (51.7%) | 13 (37.1%) | 12 (38.7%) | 62 (35.6%) | |
| wild_type | 14 (82.4%) | 43 (69.4%) | 14 (48.3%) | 22 (62.9%) | 19 (61.3%) | 112 (64.4%) | |
| **BRAF Status** | | | | | | | < 0.001 |
| N-Miss | 6 | 33 | 14 | 17 | 19 | 89 | |

| | CMS1 (N=20) | CMS2 (N=77) | CMS3 (N=35) | CMS4 (N=47) | UNK (N=43) | Total (N=222) | p value |
|---|---|---|---|---|---|---|---|
| mutant | 8 (57.1%) | 0 (0.0%) | 2 (9.5%) | 1 (3.3%) | 4 (16.7%) | 15 (11.3%) | |
| wild_type | 6 (42.9%) | 44 (100.0%) | 19 (90.5%) | 29 (96.7%) | 20 (83.3%) | 118 (88.7%) | |
| **P53 Status** | | | | | | | 0.015 |
| N-Miss | 6 | 33 | 14 | 17 | 19 | 89 | |
| mutant | 4 (28.6%) | 32 (72.7%) | 14 (66.7%) | 10 (33.3%) | 15 (62.5%) | 75 (56.4%) | |
| wild_type | 10 (71.4%) | 12 (27.3%) | 7 (33.3%) | 20 (66.7%) | 9 (37.5%) | 58 (43.6%) | |
| **OS** | | | | | | | 0.328 |
| N-Miss | 0 | 2 | 0 | 1 | 0 | 3 | |
| Events | 7 | 20 | 18 | 16 | 18 | 79 | |
| Median Survival | NA | 164.000 | 76.000 | NA | NA | 164.000 | |
| **DFS** | | | | | | | 0.545 |
| N-Miss | 0 | 2 | 1 | 1 | 3 | 7 | |
| Events | 4 | 15 | 8 | 14 | 11 | 52 | |
| Median Survival | NA | NA | NA | NA | NA | NA | |
| **DSS** | | | | | | | 0.545 |
| N-Miss | 15 | 50 | 22 | 37 | 29 | 153 | |
| Events | 0 | 1 | 1 | 0 | 2 | 4 | |
| Median Survival | NA | NA | NA | NA | NA | NA | |
| **Bulk Transcriptomics Technology (used For CMS prediction)** | | | | | | | 0.251 |
| Microarray | 3 (15.0%) | 15 (19.5%) | 9 (25.7%) | 14 (29.8%) | 17 (39.5%) | 58 (26.1%) | |
| RNA | 17 (85.0%) | 62 (80.5%) | 26 (74.3%) | 33 (70.2%) | 26 (60.5%) | 164 (73.9%) | |
| **TMA Cores (used for CellDive)** | | | | | | | 0.084 |
| Mean (SD) | 2.800 (0.523) | 2.909 (0.492) | 2.943 (0.338) | 3.000 (0.000) | 3.070 (0.632) | 2.955 (0.454) | |
| Median | 3.000 | 3.000 | 3.000 | 3.000 | 3.000 | 3.000 | |
| Q1, Q3 | 3.000, 3.000 | 3.000, 3.000 | 3.000, 3.000 | 3.000, 3.000 | 3.000, 3.000 | 3.000, 3.000 | |
| Range | 1.000 - 3.000 | 2.000 - 6.000 | 1.000 - 3.000 | 3.000 - 3.000 | 1.000 - 6.000 | 1.000 - 6.000 | |
| **Cohort** | | | | | | | 0.545 |
| Rcsi | 5 (25.0%) | 28 (36.4%) | 13 (37.1%) | 11 (23.4%) | 14 (32.6%) | 71 (32.0%) | |
| Taxonomy | 15 (75.0%) | 49 (63.6%) | 22 (62.9%) | 36 (76.6%) | 29 (67.4%) | 151 (68.0%) | |

### Table 2 Cell Count Summary

| | CMS1 (N=52) | CMS2 (N=217) | CMS3 (N=96) | CMS4 (N=136) | UNK (N=131) | Total (N=632) | p value |
|---|---|---|---|---|---|---|---|
| **Cancer Cells Count** | | | | | | | < 0.001 |
| Mean (SD) | 1826.615 (857.250) | 1854.811 (725.948) | 1597.010 (688.753) | 1527.824 (597.766) | 1627.382 (769.454) | 1695.826 (726.832) | |
| Range | 400.000 - 3524.000 | 195.000 - 3704.000 | 161.000 - 3575.000 | 275.000 - 3553.000 | 149.000 - 4135.000 | 149.000 - 4135.000 | |
| **Stromal Cells Count** | | | | | | | 0.002 |
| Mean (SD) | 1002.096 (388.939) | 1058.088 (382.520) | 1103.750 (392.227) | 1129.434 (422.881) | 1245.267 (476.359) | 1114.568 (419.806) | |
| Range | 385.000 - 2327.000 | 347.000 - 3575.000 | 379.000 - 2064.000 | 341.000 - 2620.000 | 305.000 - 2993.000 | 305.000 - 3575.000 | |
| **Immune Cells Count** | | | | | | | 0.044 |
| N-Miss | 0 | 5 | 1 | 3 | 5 | 14 | |
| Mean (SD) | 780.500 (785.377) | 438.358 (351.214) | 375.221 (352.096) | 397.602 (373.119) | 413.087 (362.723) | 443.518 (423.598) | |
| Range | 28.000 - 3109.000 | 1.000 - 1767.000 | 4.000 - 1675.000 | 1.000 - 1995.000 | 20.000 - 2076.000 | 1.000 - 3109.000 | |
| **Immune/Cancer Ratio** | | | | | | | 0.479 |
| N-Miss | 0 | 5 | 1 | 3 | 5 | 14 | |
| Mean (SD) | 0.691 (1.049) | 0.334 (0.474) | 0.345 (0.631) | 0.361 (0.666) | 0.450 (1.284) | 0.395 (0.817) | |
| Range | 0.008 - 4.013 | 0.001 - 3.732 | 0.003 - 5.438 | 0.001 - 5.938 | 0.013 - 13.403 | 0.001 - 13.403 | |
| **CD3+** | | | | | | | 0.005 |
| Mean (SD) | 530.923 (576.577) | 253.507 (242.535) | 233.135 (255.821) | 229.346 (296.791) | 231.939 (204.660) | 263.568 (301.429) | |
| Range | 7.000 - 2519.000 | 0.000 - 1436.000 | 1.000 - 1159.000 | 0.000 - 1821.000 | 2.000 - 1067.000 | 0.000 - 2519.000 | |
| **CD4+** | | | | | | | 0.014 |
| Mean (SD) | 400.404 (430.685) | 330.687 (257.679) | 239.375 (213.036) | 260.294 (185.553) | 278.756 (213.568) | 296.641 (251.899) | |
| Range | 17.000 - 1728.000 | 0.000 - 1327.000 | 7.000 - 1067.000 | 1.000 - 913.000 | 8.000 - 1052.000 | 0.000 - 1728.000 | |
| **CD8+** | | | | | | | < 0.001 |
| Mean (SD) | 327.558 (391.237) | 83.194 (93.075) | 105.531 (150.569) | 123.699 (264.967) | 99.183 (155.814) | 118.723 (207.152) | |
| Range | 1.000 - 1451.000 | 0.000 - 554.000 | 0.000 - 952.000 | 0.000 - 1762.000 | 1.000 - 1250.000 | 0.000 - 1762.000 | |
| **CD20+** | | | | | | | 0.329 |
| Mean (SD) | 92.923 (160.786) | 49.917 (83.576) | 61.135 (104.474) | 46.522 (76.819) | 77.771 (169.439) | 60.203 (116.320) | |
| Range | 0.000 - 633.000 | 0.000 - 567.000 | 0.000 - 549.000 | 0.000 - 433.000 | 0.000 - 1131.000 | 0.000 - 1131.000 | |
| **FOXP3+** | | | | | | | 0.354 |
| Mean (SD) | 164.442 (342.717) | 121.037 (215.016) | 98.792 (130.541) | 118.029 (297.999) | 103.084 (187.607) | 116.861 (233.274) | |
| Range | 1.000 - 2110.000 | 0.000 - 2225.000 | 0.000 - 647.000 | 0.000 - 2826.000 | 0.000 - 1728.000 | 0.000 - 2826.000 | |

| | CMS1 (N=52) | CMS2 (N=217) | CMS3 (N=96) | CMS4 (N=136) | UNK (N=131) | Total (N=632) | p value |
|---|---|---|---|---|---|---|---|
| **PD1+** | | | | | | | 0.014 |
| Mean (SD) | 208.481 (292.794) | 62.691 (75.366) | 111.927 (185.301) | 78.169 (109.034) | 84.466 (160.375) | 90.009 (152.997) | |
| Range | 0.000 - 1332.000 | 0.000 - 465.000 | 0.000 - 1416.000 | 0.000 - 649.000 | 0.000 - 1283.000 | 0.000 - 1416.000 | |
| **B Cell Counts** | | | | | | | 0.960 |
| N-Miss | 3 | 27 | 4 | 25 | 15 | 74 | |
| Mean (SD) | 95.735 (158.004) | 53.089 (84.289) | 60.543 (103.179) | 53.667 (78.586) | 80.362 (168.331) | 63.848 (116.531) | |
| Range | 1.000 - 594.000 | 1.000 - 560.000 | 1.000 - 549.000 | 1.000 - 433.000 | 1.000 - 1131.000 | 1.000 - 1131.000 | |
| **Cytotoxic T Cell Counts** | | | | | | | 0.002 |
| N-Miss | 0 | 10 | 6 | 16 | 5 | 37 | |
| Mean (SD) | 241.500 (290.354) | 64.232 (71.012) | 86.344 (117.757) | 110.058 (239.098) | 68.627 (92.481) | 93.242 (163.486) | |
| Range | 1.000 - 1011.000 | 1.000 - 418.000 | 1.000 - 668.000 | 1.000 - 1602.000 | 1.000 - 543.000 | 1.000 - 1602.000 | |
| **Helper T Cell Counts** | | | | | | | 0.036 |
| N-Miss | 1 | 10 | 2 | 8 | 5 | 26 | |
| Mean (SD) | 133.216 (154.050) | 111.594 (114.295) | 82.926 (100.644) | 74.023 (76.625) | 93.516 (106.817) | 97.272 (109.017) | |
| Range | 2.000 - 650.000 | 1.000 - 635.000 | 1.000 - 629.000 | 1.000 - 536.000 | 1.000 - 543.000 | 1.000 - 650.000 | |
| **Regulatory T Cell Counts** | | | | | | | 0.010 |
| N-Miss | 4 | 9 | 9 | 14 | 13 | 49 | |
| Mean (SD) | 57.583 (78.932) | 46.438 (53.459) | 35.230 (52.441) | 34.148 (37.979) | 36.271 (36.879) | 41.053 (50.516) | |
| Range | 1.000 - 416.000 | 1.000 - 317.000 | 1.000 - 306.000 | 1.000 - 227.000 | 1.000 - 180.000 | 1.000 - 416.000 | |
| **DN T Cell Counts** | | | | | | | < 0.001 |
| N-Miss | 4 | 21 | 9 | 16 | 10 | 60 | |
| Mean (SD) | 106.125 (205.412) | 30.582 (66.684) | 25.540 (49.311) | 25.183 (49.931) | 26.876 (34.015) | 34.238 (81.240) | |
| Range | 1.000 - 977.000 | 1.000 - 690.000 | 1.000 - 308.000 | 1.000 - 434.000 | 1.000 - 236.000 | 1.000 - 977.000 | |
| **CD3- CD4+ or CD8+ Cell Counts** | | | | | | | 0.001 |
| N-Miss | 0 | 8 | 1 | 4 | 5 | 18 | |
| Mean (SD) | 167.019 (207.859) | 147.349 (113.575) | 97.084 (80.139) | 129.197 (88.413) | 117.183 (90.185) | 131.145 (112.715) | |
| Range | 3.000 - 1034.000 | 2.000 - 880.000 | 2.000 - 443.000 | 1.000 - 504.000 | 3.000 - 531.000 | 1.000 - 1034.000 | |

### Table 3 Cell Network Scores Summary

| | CMS1 (N=52) | CMS2 (N=217) | CMS3 (N=96) | CMS4 (N=136) | UNK (N=131) | Total (N=632) | p value |
|---|---|---|---|---|---|---|---|
| **Average Shortest Distance from Cytotoxic T cell to Cancer Cell** | | | | | | | 0.005 |
| N-Miss | 0 | 10 | 6 | 16 | 5 | 37 | |
| Mean (SD) | 74.166 (56.261) | 103.300 (82.848) | 100.607 (66.300) | 102.896 (67.172) | 103.385 (83.326) | 100.283 (75.777) | |
| Range | 27.411 - 320.187 | 9.220 - 583.369 | 25.951 - 313.096 | 14.036 - 392.792 | 27.203 - 646.699 | 9.220 - 646.699 | |
| **Average Shortest Distance from Regulatory T cell to Cancer Cell** | | | | | | | 0.057 |
| N-Miss | 4 | 9 | 9 | 14 | 13 | 49 | |
| Mean (SD) | 97.050 (65.593) | 117.051 (77.239) | 116.739 (65.852) | 121.399 (80.063) | 125.159 (80.404) | 117.909 (76.105) | |
| Range | 32.785 - 338.285 | 29.887 - 502.255 | 29.340 - 376.390 | 18.028 - 576.880 | 26.378 - 489.964 | 18.028 - 576.880 | |
| **Average Shortest Distance from Helper T cell to Cancer Cell** | | | | | | | 0.008 |
| N-Miss | 1 | 10 | 2 | 8 | 5 | 26 | |
| Mean (SD) | 89.163 (57.847) | 116.283 (82.004) | 108.692 (64.797) | 119.928 (76.189) | 122.415 (91.429) | 114.868 (78.974) | |
| Range | 33.571 - 248.986 | 14.422 - 591.021 | 27.203 - 444.974 | 26.249 - 468.337 | 28.838 - 762.222 | 14.422 - 762.222 | |
| **Average Shortest Distance from B cell to Cancer Cell** | | | | | | | 0.007 |
| N-Miss | 3 | 27 | 4 | 25 | 15 | 74 | |
| Mean (SD) | 84.780 (59.500) | 103.392 (83.400) | 87.070 (64.148) | 112.893 (86.759) | 121.205 (107.503) | 104.659 (85.832) | |
| Range | 20.616 - 251.571 | 19.542 - 550.343 | 10.296 - 417.366 | 26.215 - 592.851 | 23.543 - 796.883 | 10.296 - 796.883 | |
| **Average Shortest Distance from DN T cell to Stromal Cell** | | | | | | | 0.007 |
| N-Miss | 4 | 21 | 9 | 16 | 10 | 60 | |
| Mean (SD) | 54.639 (20.862) | 47.013 (16.463) | 44.075 (12.906) | 44.493 (15.066) | 47.649 (20.481) | 46.812 (17.219) | |
| Range | 27.459 - 141.689 | 12.042 - 129.403 | 24.885 - 84.737 | 22.091 - 98.250 | 21.190 - 164.639 | 12.042 - 164.639 | |
| **Average #No Cancer Cells near Cytotoxic T Cells** | | | | | | | 0.005 |
| N-Miss | 0 | 10 | 6 | 16 | 5 | 37 | |
| Mean (SD) | 13.930 (7.772) | 10.573 (6.924) | 9.606 (6.587) | 9.473 (6.920) | 9.963 (6.632) | 10.369 (6.969) | |
| Range | 3.075 - 37.000 | 0.000 - 31.103 | 0.000 - 28.600 | 0.667 - 34.000 | 0.000 - 24.938 | 0.000 - 37.000 | |
| **Average #No Cancer Cells near Regulatory T Cells** | | | | | | | 0.001 |
| N-Miss | 4 | 9 | 9 | 14 | 13 | 49 | |
| Mean (SD) | 9.398 (6.859) | 7.747 (5.509) | 6.619 (5.958) | 6.470 (5.646) | 5.890 (4.672) | 7.071 (5.644) | |

| | CMS1 (N=52) | CMS2 (N=217) | CMS3 (N=96) | CMS4 (N=136) | UNK (N=131) | Total (N=632) | p value |
|---|---|---|---|---|---|---|---|
| Range | 0.000 - 22.667 | 0.000 - 28.650 | 0.000 - 28.545 | 0.000 - 26.600 | 0.000 - 23.000 | 0.000 - 28.650 | |
| **Average #No Cancer Cells near Helper T Cells** | | | | | | | 0.006 |
| N-Miss | 1 | 10 | 2 | 8 | 5 | 26 | |
| Mean (SD) | 10.367 (6.319) | 7.949 (5.513) | 7.695 (5.919) | 6.882 (5.324) | 7.283 (5.675) | 7.749 (5.695) | |
| Range | 0.805 - 25.750 | 0.000 - 29.154 | 0.000 - 31.667 | 0.000 - 24.167 | 0.000 - 29.000 | 0.000 - 31.667 | |
| **Average #No Cancer Cells near DN T Cells** | | | | | | | 0.010 |
| N-Miss | 4 | 21 | 9 | 16 | 10 | 60 | |
| Mean (SD) | 13.937 (6.185) | 12.856 (7.553) | 11.370 (7.115) | 10.644 (7.216) | 11.154 (7.190) | 11.897 (7.289) | |
| Range | 4.000 - 29.000 | 0.000 - 34.800 | 0.200 - 35.833 | 0.000 - 28.333 | 0.000 - 35.538 | 0.000 - 35.833 | |
| **Average #No Stromal Cells near Cytotoxic T Cells** | | | | | | | < 0.001 |
| N-Miss | 0 | 10 | 6 | 16 | 5 | 37 | |
| Mean (SD) | 8.126 (3.746) | 10.404 (3.577) | 11.432 (4.102) | 10.815 (4.303) | 11.209 (5.101) | 10.614 (4.253) | |
| Range | 2.000 - 15.667 | 2.150 - 22.000 | 2.989 - 22.000 | 2.800 - 27.304 | 2.375 - 24.846 | 2.000 - 27.304 | |
| **Average #No Stromal Cells near Regulatory T Cells** | | | | | | | 0.005 |
| N-Miss | 4 | 9 | 9 | 14 | 13 | 49 | |
| Mean (SD) | 9.389 (4.730) | 11.786 (3.870) | 12.537 (5.384) | 12.350 (5.084) | 12.722 (5.349) | 12.008 (4.826) | |
| Range | 0.850 - 21.333 | 1.273 - 27.000 | 2.000 - 38.000 | 4.000 - 31.000 | 2.214 - 27.909 | 0.850 - 38.000 | |
| **Average #No Stromal Cells near Helper T Cells** | | | | | | | < 0.001 |
| N-Miss | 1 | 10 | 2 | 8 | 5 | 26 | |
| Mean (SD) | 9.252 (4.215) | 11.550 (3.535) | 12.447 (4.101) | 12.312 (4.369) | 12.239 (4.914) | 11.800 (4.246) | |
| Range | 1.388 - 19.839 | 2.391 - 19.583 | 3.674 - 21.900 | 3.500 - 28.042 | 2.125 - 23.614 | 1.388 - 28.042 | |
| **Average #No Stromal Cells near DN T Cells** | | | | | | | 0.022 |
| N-Miss | 4 | 21 | 9 | 16 | 10 | 60 | |
| Mean (SD) | 8.110 (3.484) | 10.128 (4.181) | 10.689 (4.449) | 10.280 (4.789) | 10.746 (5.758) | 10.207 (4.708) | |
| Range | 1.841 - 16.000 | 1.000 - 23.250 | 3.000 - 19.500 | 0.667 - 23.600 | 0.000 - 26.667 | 0.000 - 26.667 | |
| **Pct of Regulatory T cells inside Stromal Cell Clusters** | | | | | | | < 0.001 |
| N-Miss | 5 | 9 | 9 | 15 | 14 | 52 | |
| Mean (SD) | 0.830 (0.179) | 0.888 (0.117) | 0.917 (0.121) | 0.918 (0.105) | 0.908 (0.101) | 0.898 (0.120) | |
| Range | 0.300 - 1.000 | 0.165 - 1.000 | 0.333 - 1.000 | 0.333 - 1.000 | 0.600 - 1.000 | 0.165 - 1.000 | |

| | CMS1 (N=52) | CMS2 (N=217) | CMS3 (N=96) | CMS4 (N=136) | UNK (N=131) | Total (N=632) | p value |
|---|---|---|---|---|---|---|---|
| **Degree Centrality Stromal Cells** | | | | | | | < 0.001 |
| Mean (SD) | 0.588 (0.171) | 0.613 (0.134) | 0.674 (0.138) | 0.667 (0.136) | 0.656 (0.142) | 0.641 (0.143) | |
| Range | 0.179 - 0.895 | 0.311 - 0.955 | 0.341 - 0.994 | 0.368 - 0.968 | 0.217 - 0.979 | 0.179 - 0.994 | |
| **Average Clustering Coefficient Cancer Cells** | | | | | | | < 0.001 |
| Mean (SD) | 0.507 (0.026) | 0.514 (0.018) | 0.524 (0.020) | 0.516 (0.022) | 0.512 (0.026) | 0.515 (0.022) | |
| Range | 0.460 - 0.559 | 0.477 - 0.580 | 0.472 - 0.581 | 0.469 - 0.576 | 0.450 - 0.610 | 0.450 - 0.610 | |
| **Cancer - Immune Neighbourhood Enrichment** | | | | | | | < 0.001 |
| N-Miss | 0 | 5 | 1 | 3 | 5 | 14 | |
| Mean (SD) | -22.801 (14.755) | -32.151 (14.459) | -24.251 (14.392) | -25.891 (13.340) | -25.035 (13.244) | -27.352 (14.399) | |
| Range | -57.883 - 6.030 | -69.173 - 4.882 | -67.646 - 1.824 | -59.685 - 0.903 | -66.128 - 1.268 | -69.173 - 6.030 | |
| **Cancer - Stroma Neighbourhood Enrichment** | | | | | | | < 0.001 |
| Mean (SD) | -29.239 (13.016) | -37.239 (12.086) | -32.746 (11.258) | -34.613 (12.500) | -36.895 (12.971) | -35.262 (12.522) | |
| Range | -55.436 - -0.729 | -68.593 - -2.294 | -62.139 - -0.485 | -60.329 - -4.623 | -66.975 - 2.981 | -68.593 - 2.981 | |
| **Stroma - Immune Neighbourhood Enrichment** | | | | | | | < 0.001 |
| N-Miss | 0 | 5 | 1 | 3 | 5 | 14 | |
| Mean (SD) | -3.453 (12.764) | 7.157 (11.544) | 3.636 (11.431) | 4.877 (9.564) | 2.797 (12.150) | 4.343 (11.700) | |
| Range | -35.114 - 19.859 | -39.576 - 28.747 | -33.035 - 28.013 | -22.019 - 25.305 | -51.813 - 22.847 | -51.813 - 28.747 | |
| **Immune - Immune Neighbourhood Enrichment** | | | | | | | 0.001 |
| N-Miss | 0 | 6 | 1 | 4 | 5 | 16 | |
| Mean (SD) | 26.598 (14.627) | 30.140 (13.528) | 25.046 (14.071) | 25.438 (12.466) | 25.760 (14.202) | 27.152 (13.762) | |
| Range | 2.540 - 55.360 | 2.867 - 78.426 | 2.646 - 66.894 | -0.372 - 66.658 | -0.292 - 71.386 | -0.372 - 78.426 | |
| **Cancer - Cancer Neighbourhood Enrichment** | | | | | | | 0.003 |
| Mean (SD) | 45.104 (14.543) | 52.532 (15.060) | 46.912 (14.063) | 49.246 (14.109) | 51.162 (15.079) | 50.076 (14.831) | |
| Range | 7.598 - 81.597 | 17.714 - 94.110 | 20.494 - 89.881 | 17.385 - 77.076 | 12.532 - 91.312 | 7.598 - 94.110 | |
